# Superior Frontal Gyrus Cortical Thickness Mediates the Association Between Openness to Experience and Theory of Mind: Evidence from the Human Connectome Project

**DOI:** 10.64898/2026.10.31.763281

**Authors:** Barış Akar, Ümit Atasever

## Abstract

**Background:** Theory of mind (ToM), the capacity to infer the mental states of others, varies considerably across healthy adults. Openness to experience has been linked both to social-cognitive abilities and to thinner prefrontal cortex, but whether cortical structure statistically accounts for the association between openness and ToM is unknown. We tested whether bilateral superior frontal gyrus (SFG) cortical thickness mediates this association.

**Methods:** Data from 1,049 healthy young adults with complete data were drawn from the Human Connectome Project. ToM was indexed by accuracy on the Heider–Simmel social animation task, openness by the NEO Five-Factor Inventory, and SFG thickness by FreeSurfer. Models adjusted for age, sex, intracranial volume and fluid intelligence; indirect effects were tested with 5,000 bootstrap resamples, and Benjamini–Hochberg false discovery rate correction was applied.

**Results:** Openness showed a positive total effect on ToM accuracy (β = .081, *p* = .009). Higher openness was associated with thinner left and right SFG (β = −.085 and −.084), and thinner SFG with higher ToM accuracy (β = −.082 and −.075). Indirect effects were significant (left: .0069, 95% CI [.0010, .0158]; right: .0063, 95% CI [.0007, .0146]), accounting for approximately 8.5% and 7.8% of the total effect, and remained significant after rank-based normalisation of the ceiling-affected ToM scores.

**Conclusions:** SFG cortical thickness partially and statistically mediates the association between openness to experience and social cognition. Because the data are cross-sectional and effects are small, the findings do not establish a causal mechanism, neural efficiency or synaptic pruning, but they identify a candidate structural pathway for longitudinal testing.

## 1. Introduction

Theory of mind (ToM)—the ability to attribute beliefs, intentions and emotions to others—is a cornerstone of human social life.^1^ Classic false-belief paradigms showed that this capacity typically emerges in early childhood and is selectively affected in autism,^2^ and neuroimaging meta-analyses have identified a reliable mentalising network comprising the medial prefrontal cortex, temporoparietal junction, precuneus and temporal poles.^3–5^ Even among healthy adults, however, ToM performance varies, and the dispositional and neuroanatomical factors underlying this variability remain incompletely understood.

Openness to experience, the Big Five trait capturing imagination, aesthetic sensitivity, intellectual curiosity and receptivity to new ideas,^6^ is a plausible contributor. Individuals high in openness readily adopt alternative perspectives and engage with ambiguous, abstract information, and openness predicts the efficiency of the default network,^7^ which overlaps substantially with the mentalising network. Personality neuroscience has also linked Big Five traits to regional brain structure.^8^ Notably, surface-based studies—including a study of the full Human Connectome Project (HCP) sample—found that higher openness is associated with thinner cortex in prefrontal regions, including the superior prefrontal cortex.^9,10^ Such associations are small and only partly reproducible across samples,^11^ emphasising the need for large datasets and theoretically constrained hypotheses.

The superior frontal gyrus (SFG) is a promising candidate for linking openness and ToM. It spans the dorsolateral and dorsomedial frontal cortex; its medial portion is functionally coupled with the default network and overlaps with dorsomedial prefrontal regions engaged during mentalising.^12,13^ Cortical thinning in frontal association cortex continues from adolescence into early adulthood,^14,15^ and thinner medial prefrontal cortex has been associated with better spontaneous mentalising in young adults.^16^ Thinner cortex in this age range may therefore reflect maturational refinement rather than deficit.

We tested whether bilateral SFG cortical thickness statistically mediates the association between openness to experience and ToM accuracy in a large sample of healthy young adults from the HCP. We hypothesised that (H1) openness would positively predict ToM; (H2) openness would be associated with thinner SFG; (H3) thinner SFG would be associated with higher ToM accuracy; and (H4) the indirect effect via SFG thickness would differ from zero (Figure 2).

**Figure 1.**
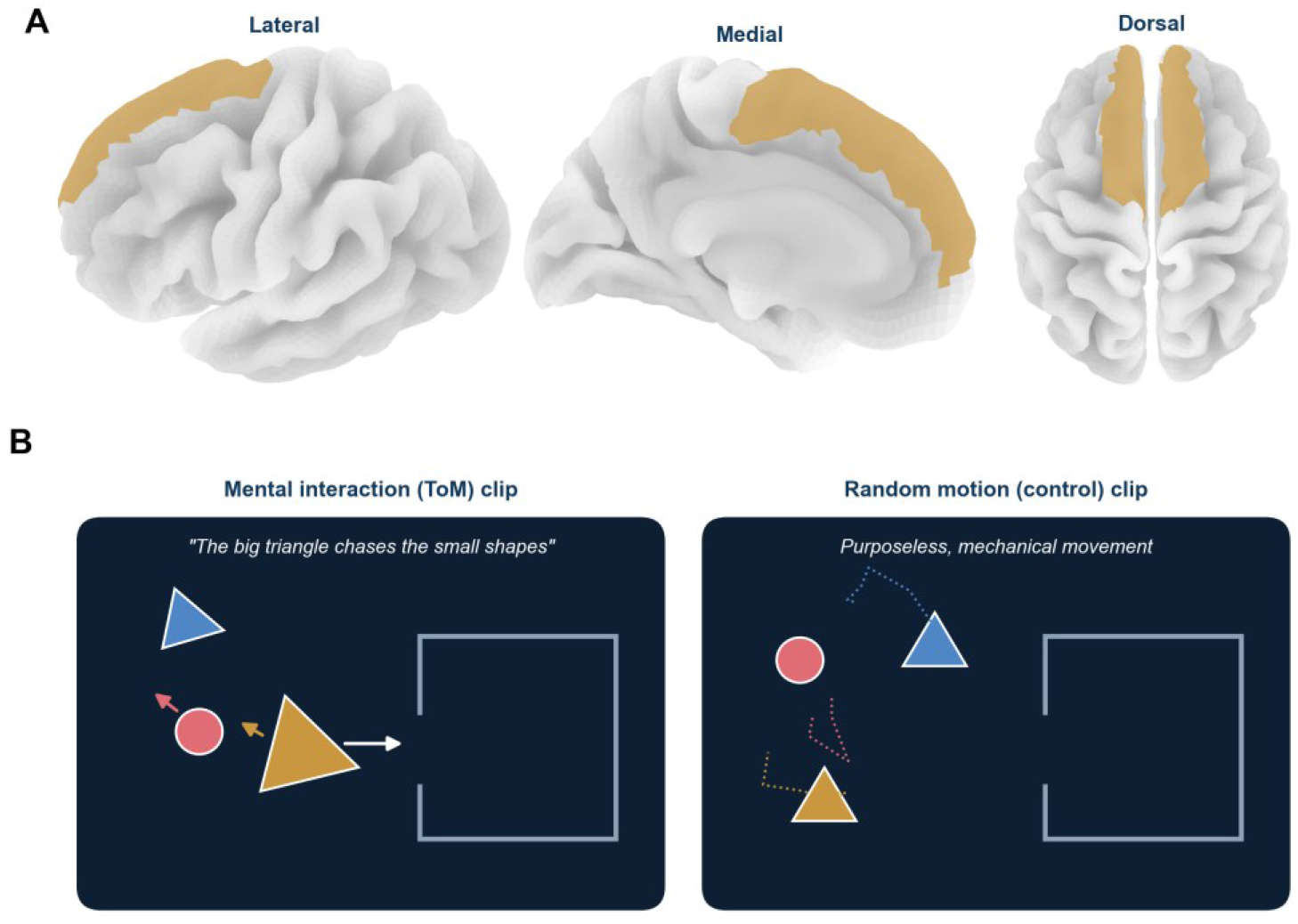
Region of interest and ToM task. (A) Superior frontal gyrus (gold) on the left-hemisphere pial surface (lateral and medial views) and on both hemispheres (dorsal view); display uses the Destrieux atlas on fsaverage, whereas analyses used the Desikan–Killiany superior frontal parcel. (B) Schematic of the Heider–Simmel social animation task: in mental-interaction clips, shapes appear to act with intentions; in control clips, they move randomly.

**Figure 2.**
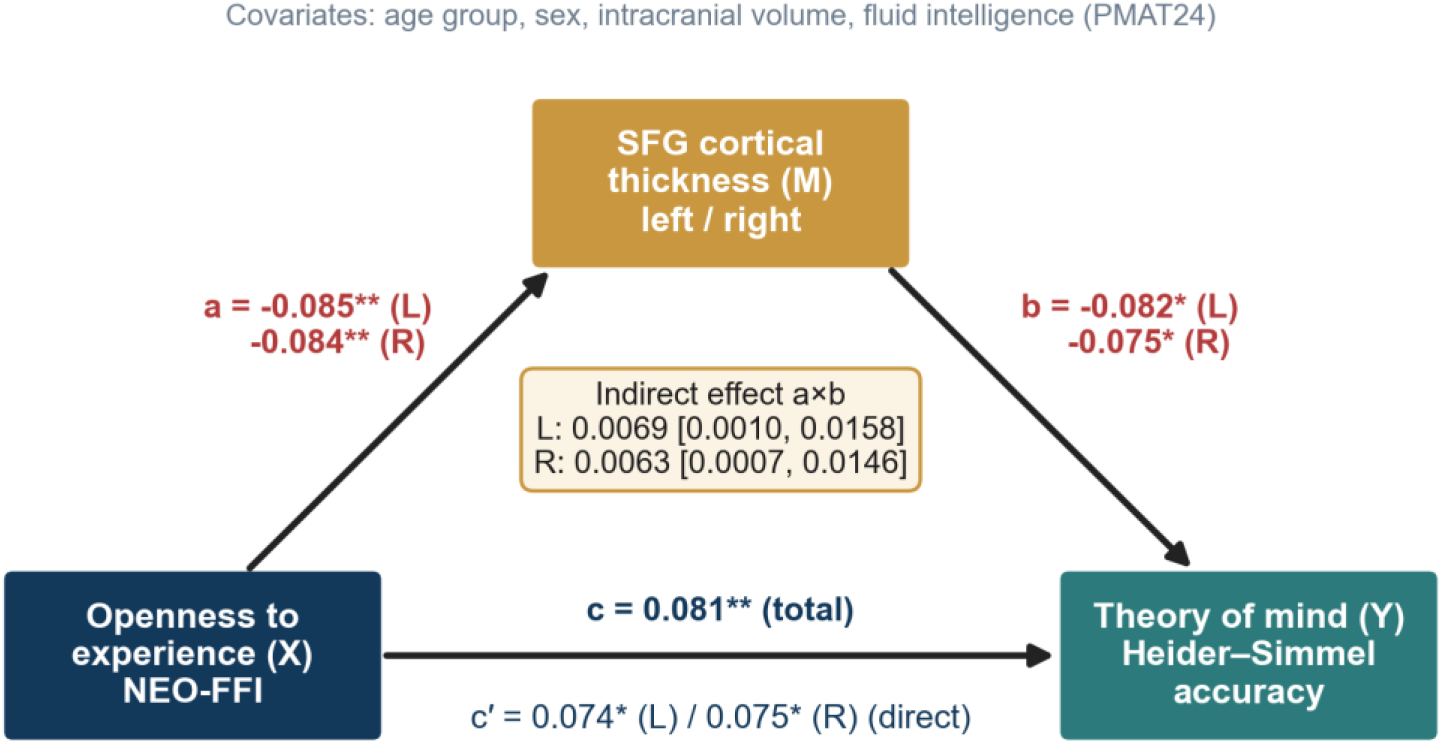
Mediation model with standardised coefficients. Values for the left (L) and right (R) SFG are shown for the *a* and *b* paths and the direct effect (*c*′); the total effect (*c*) is shared by both models. Indirect effects are given with 95% bootstrap confidence intervals (5,000 resamples). * *p* < .05, ** *p* < .01 (FDR-corrected for *a* and *b*).

## 2. Materials and Methods

### 2.1. Participants

Data were obtained from the WU-Minn HCP S1200 open-access release.^17^ Participants were healthy adults aged 22–37 years. We included all participants with complete data on ToM accuracy, openness, left and right SFG thickness, age, sex, intracranial volume and fluid intelligence, yielding a sample of 1,049 (562 women, 53.6%). In the open-access release, age is provided in bands (22–25: 21.2%; 26–30: 43.6%; 31–35: 34.2%; 36+: 1.0%).

### 2.2. Measures

#### Theory of mind

*Theory of mind*. ToM was assessed with the HCP social cognition task, adapted from the Heider–Simmel animations.^18–20^ Participants viewed 20-s video clips of moving geometric shapes that either interacted in a socially meaningful way (e.g., chasing, coaxing) or moved randomly, and judged whether a mental interaction had occurred (Figure 1B). Such animations reliably elicit mental-state attributions and recruit the mentalising network, and individuals with autism attribute fewer mental states to them.^19,21^ The dependent variable was the percentage of ToM clips correctly identified as involving mental interaction (Social_Task_TOM_Perc_TOM).

#### Openness to experience

Openness was measured with the 12-item openness scale of the NEO Five-Factor Inventory (NEO-FFI; range 0–48).^6^ *Fluid intelligence*. Fluid intelligence was indexed by the number of correct responses on the 24-item Penn Progressive Matrices (PMAT24).^22^

#### SFG cortical thickness

Structural T1-weighted images (3 T, 0.7-mm isotropic) were processed with the HCP minimal preprocessing pipelines, which incorporate FreeSurfer.^23,24^ Mean cortical thickness of the left and right superior frontal regions of the Desikan–Killiany atlas^25^ and FreeSurfer-estimated intracranial volume were taken from the HCP release. For illustration, the SFG is displayed on the fsaverage surface using the Destrieux atlas (Figure 1A).^26^

#### 2.3. Statistical analysis

Continuous variables (ToM accuracy, openness, SFG thickness, PMAT24 and intracranial volume) were z-standardised so that coefficients represent standardised effects (β). Age band (reference: 22–25 years) and sex were dummy-coded. For each hemisphere, three ordinary least squares models were estimated: (i) ToM regressed on openness and covariates (total effect, *c*); (ii) SFG thickness regressed on openness and covariates (*a* path); and (iii) ToM regressed on openness, SFG thickness and covariates (*b* path and direct effect, *c*′). The indirect effect (*a* × *b*) was tested with 5,000 nonparametric bootstrap resamples and percentile 95% confidence intervals (CIs).^27,28^ Benjamini–Hochberg false discovery rate (FDR) correction was applied across the two hemispheric models for the *a, b* and indirect paths.^29^ Because ToM accuracy showed a pronounced ceiling effect, we repeated the mediation analysis after rank-based inverse-normal transformation of ToM scores as a sensitivity analysis. Analyses were performed in Python (pandas, statsmodels, SciPy).

## 3. Results

### 3.1. Descriptive statistics

Descriptive statistics and zero-order correlations are presented in Table 1. ToM accuracy was high and negatively skewed (skewness = -2.68); 77.6% of participants achieved 100% accuracy. Openness correlated positively with ToM (*r* = 0.13) and negatively with left and right SFG thickness, and SFG thickness correlated negatively with ToM. Fluid intelligence correlated with both openness and ToM (*r* = 0.23 and 0.23), underscoring the importance of its inclusion as a covariate. Left and right SFG thickness were highly correlated (*r* = 0.85).

**Table 1.** Descriptive statistics and zero-order correlations (N = 1,049)

| **Variable** | ***M* ± *SD*** | **Range** | **1** | **2** | **3** | **4** | **5** |
|---|---|---|---|---|---|---|---|
| 1. Openness (NEO-FFI) | 28.35 ± 6.22 | 10–47 | — |  |  |  |  |
| 2. Left SFG thickness (mm) | 2.84 ± 0.13 | 2.05–3.26 | −.07 | — |  |  |  |
| 3. Right SFG thickness (mm) | 2.87 ± 0.13 | 2.41–3.29 | −.06 | .85 | — |  |  |
| 4. ToM accuracy (%) | 94.07 ± 12.65 | 0–100 | .13 | −.10 | −.08 | — |  |
| 5. Fluid intelligence (PMAT24) | 16.98 ± 4.74 | 4–24 | .23 | −.02 | .01 | .23 | — |
| Intracranial volume (cm³) | 1585 ± 187 | 831–2143 |  |  |  |  |  |

*Note*. Pearson correlations; with N > 1,000, |*r*| ≥ .07 corresponds to uncorrected *p* < .05. NEO-FFI, NEO Five-Factor Inventory; PMAT24, Penn Progressive Matrices; SFG, superior frontal gyrus; ToM, theory of mind.

### 3.2. Mediation analysis

After adjustment for age, sex, intracranial volume and fluid intelligence, openness positively predicted ToM accuracy (total effect β = .081, 95% CI [.020, .141], *p* = .009; H1 supported). Fluid intelligence was the strongest covariate (β = .196, *p* < .001). Higher openness was associated with thinner SFG in both hemispheres (left β = −.085, right β = −.084; both FDR *p* = .008; H2 supported), and, controlling for openness, thinner SFG was associated with higher ToM accuracy (left β = −.082, right β = −.075; both FDR *p* = .015; H3 supported; Table 2; Figure 2).

**Table 2.** Standardised path coefficients of the mediation models.

| **Path** | **Left β** | **95% CI** | ***p*** | **Right β** | **95% CI** | ***p*** |
|---|---|---|---|---|---|---|
| Total effect, c (X → Y) | .081 | [.020, .141] | .009 | .081 | [.020, .141] | .009 |
| a path (X → M) | −.085 | [−.146, −.023] | .008 | −.084 | [−.145, −.022] | .008 |
| b path (M → Y \| X) | −.082 | [−.142, −.021] | .015 | −.075 | [−.135, −.015] | .015 |
| Direct effect, c′ (X → Y \| M) | .074 | [.013, .135] | .017 | .075 | [.014, .135] | .016 |
| Indirect effect, a × b (bootstrap) | .0069 | [.0010, .0158] | .019 | .0063 | [.0007, .0146] | .019 |
| Proportion mediated | 8.5% |  |  | 7.8% |  |  |
| *Sensitivity: rank-normalised ToM* |  |  |  |  |  |  |
| Indirect effect, a × b | .0063 | [.0004, .0148] |  | .0056 | [.0002, .0133] |  |

*Note.* All models adjusted for age band, sex, intracranial volume and fluid intelligence (N = 1,049). *p* values for the *a*, *b* and indirect paths are Benjamini–Hochberg FDR-corrected across hemispheres; CIs for the indirect effect are bootstrap percentile intervals (5,000 resamples). The sensitivity analysis repeated the bootstrap mediation after rank-based inverse-normal transformation of ToM accuracy.

The product of these two negative paths yielded positive indirect effects whose bootstrap CIs excluded zero (left: a × b = .0069, 95% CI [.0010, .0158]; right: .0063, 95% CI [.0007, .0146]; FDR *p* = .019; H4 supported; Figure 3). Approximately 8.5% (left) and 7.8% (right) of the total effect was transmitted via SFG thickness. Direct effects remained significant (left *c*′ = .074, right *c*′ = .075), indicating partial mediation. The full models explained approximately 7.2% of the variance in ToM accuracy. Raw-data trends across quintiles were consistent with all three paths (Figure 4).

**Figure 3.**
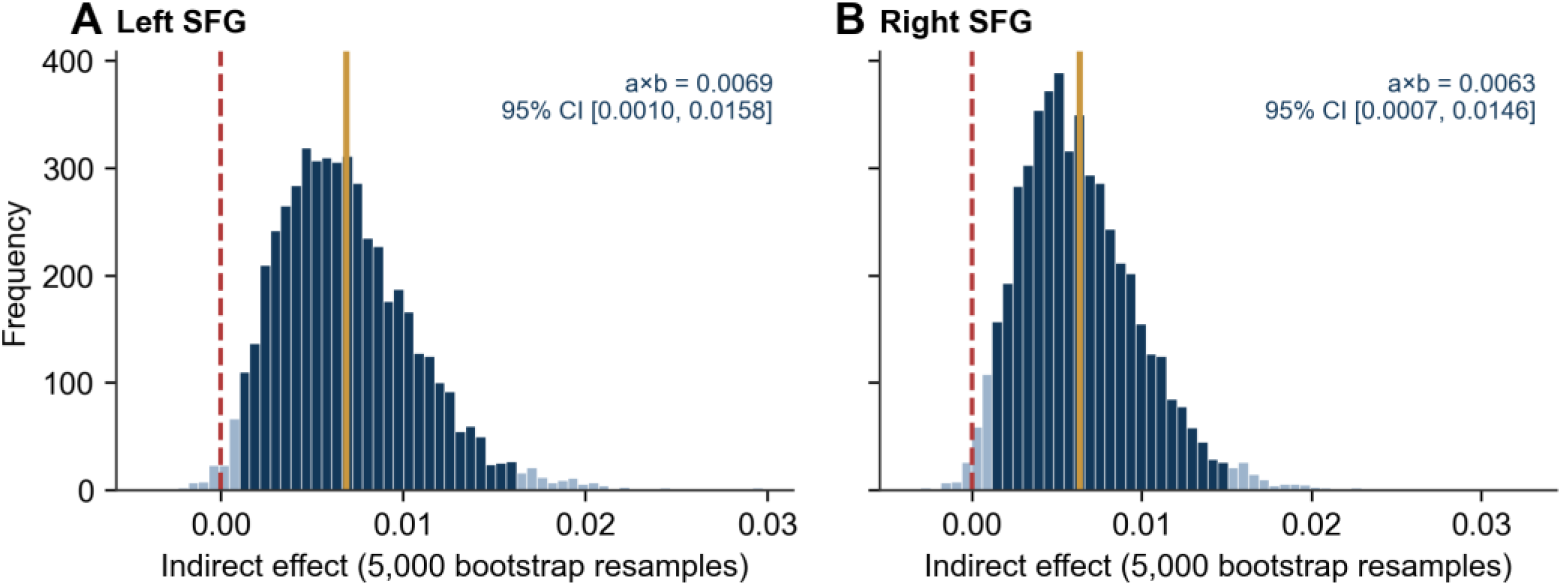
Bootstrap distributions of the indirect effects for the (A) left and (B) right SFG. Dark bars denote the 95% percentile confidence interval, the gold line the mean indirect effect, and the dashed red line zero.

**Figure 4.**
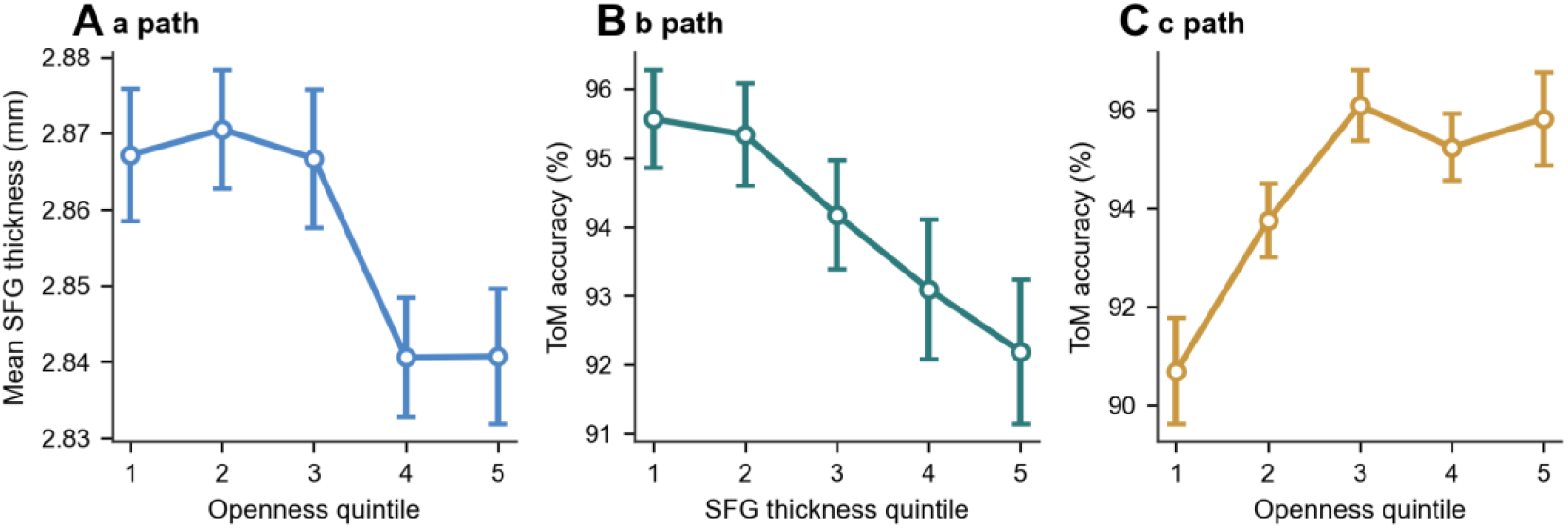
Raw-data illustration of the mediation paths. Mean ± standard error by quintile of the predictor: (A) SFG thickness (mean of hemispheres) by openness quintile; (B) ToM accuracy by SFG thickness quintile; (C) ToM accuracy by openness quintile. Values are unadjusted for covariates.

### 3.3. Sensitivity analysis

Because most participants performed at ceiling, we repeated the analysis with rank-normalised ToM scores. The total effect of openness (β = .086) and the indirect effects via left (.0063, 95% CI [.0004, .0148]) and right SFG (.0056, 95% CI [.0002, .0133]) were essentially unchanged, indicating that the results were not driven by the skewed score distribution.

## 4. Discussion

In more than one thousand healthy young adults, openness to experience was positively associated with ToM accuracy, and a small but statistically reliable portion of this association was carried by bilateral SFG cortical thickness. More open individuals had thinner SFG, and thinner SFG predicted better identification of mental interactions, yielding a positive indirect effect that was consistent across hemispheres and robust to the ceiling effect in the ToM measure.

The *a* path replicates prior reports that openness is related to thinner prefrontal cortex,^9,10^ here extended with adjustment for fluid intelligence and intracranial volume. The *b* path accords with evidence that thinner medial prefrontal cortex accompanies superior spontaneous mentalising in young adults.^16^ Together, these results suggest that the SFG—whose medial portion belongs to the dorsomedial prefrontal component of the mentalising and default networks^3,12^—may be a structural meeting point of personality and social cognition. This dovetails with findings that openness predicts the global efficiency of default network nodes.^7^

Why might thinner cortex accompany better performance? In early adulthood, frontal cortical thinning reflects ongoing maturational processes, including synaptic refinement and increasing intracortical myelination, that are thought to optimise circuit function.^14,15^ The neural efficiency hypothesis similarly proposes that more capable individuals require fewer neural resources to perform a task; for example, individuals with higher reasoning scores showed lower cortical glucose metabolism during task performance.^30^ Openness, by fostering engagement with novel and ambiguous social information, might be associated with experience-dependent refinement of frontal circuitry. We stress, however, that our cross-sectional thickness data cannot demonstrate pruning, myelination or efficiency; these interpretations remain hypotheses.

The effect sizes were small: the total effect was about 0.08 standard deviations, and SFG thickness accounted for roughly one-twelfth of it. Such magnitudes are typical of brain–behaviour associations for individual differences and require large samples for reliable detection.^31,32^ The present results should therefore be viewed as identifying a candidate pathway rather than a principal mechanism.

### 4.1. Limitations

First, the cross-sectional design precludes causal or temporal inference; statistical mediation does not imply that openness changes cortical thickness or that thickness changes ToM. Second, the HCP sample contains many twins and siblings, but family structure is available only in restricted-access data and was not modelled, which may underestimate standard errors. Third, the ToM measure showed a strong ceiling effect; although rank-based sensitivity analyses yielded similar results, measures with greater dynamic range would improve sensitivity. Fourth, the Desikan–Killiany superior frontal parcel is large and spans functionally distinct subregions; finer parcellations could localise the effect more precisely. Finally, age was available only in bands in the open-access data.

## 5. Conclusions

Openness to experience is positively associated with theory of mind in healthy young adults, and bilateral SFG cortical thickness partially mediates this association. These findings point to the SFG as a structural correlate linking personality and social cognition, while longitudinal and multimodal studies are needed to establish directionality and mechanism.

## Declarations

### Ethics statement

This study is a secondary analysis of de-identified, open-access data. HCP data collection was approved by the Washington University in St. Louis institutional review board, and all participants provided informed consent. The authors complied with the HCP Open Access Data Use Terms.

### Data availability

HCP data are available at https://db.humanconnectome.org/ under the HCP data use terms. Analysis code is available from the corresponding author upon reasonable request.

## Acknowledgements

Data were provided by the Human Connectome Project, WU-Minn Consortium (Principal Investigators: David Van Essen and Kamil Ugurbil; 1U54MH091657) funded by the 16 NIH Institutes and Centers that support the NIH Blueprint for Neuroscience Research; and by the McDonnell Center for Systems Neuroscience at Washington University. Preliminary findings were presented at the 9th BILSEL International Harput Scientific Research Congress (17–18 October 2026, Elaziğ, Türkiye).

## Funding

This research received no specific grant from any funding agency in the public, commercial or not-for-profit sectors.

## Conflict of interest

The authors declare no conflicts of interest.

## Author contributions (CRediT)

Bariş Akar: Conceptualization, Methodology, Software, Formal analysis, Visualization, Writing – original draft. Ümit Atasever: Supervision, Validation, Writing – review & editing.

## Use of generative AI

A generative AI tool was used to assist with drafting the text and producing figures from the authors’ analysis outputs. The authors reviewed and verified all content and take full responsibility for the manuscript.

## Notes

https://db.humanconnectome.org

### Competing Interest Statement

The authors have declared no competing interest.

